# Advancing Neurological Liquid Biopsy via Six-Base Sequencing of Cerebrospinal Fluid Cell-Free DNA

**DOI:** 10.64898/2026.09.06.749692

**Authors:** Chiara Starvaggi Cucuzza, Chandana Rao Prakash, Cillian Nolan, Mohammad Hossein Sepehri, Angela Simeone, Luke Sarre, Tom Charlesworth, Fabio Puddu, Yanan Han, Mohsen Khademi, Virginija Danylaitė Karrenbauer, Björn Evertsson, Katharina Fink, Per Svenningsson, Caroline Ingre, Christina Seitz, Fang Fang, Radu Constantinescu, Lenka Novakova, Eric P Thelin, Fredrik Piehl, Lara Kular, Maria Needhamsen, Maja Jagodic

## Abstract

Real-time molecular insights into the human brain are crucial for understanding neurological conditions and development of soluble biomarkers but challenged by the limited access to target tissue. Here we utilize 6-base sequencing to profile DNA modifications of cell-free DNA released from neural cells into cerebrospinal fluid. This strategy provides molecular, quantitative and qualitative insights into the ongoing neurodegenerative biology, which paves the way for liquid biopsy approaches to study neurological conditions.

---

The use of soluble biomarkers in the diagnosis and monitoring of neurological diseases is still limited in clinical practice^1^. Currently approved biomarkers include cerebrospinal fluid (CSF) levels of proteins undergoing misfolding and causing neurodegeneration in Alzheimer’s disease, such as amyloid beta and phosphorylated tau, which are tightly connected to the underlying disease mechanisms. On the other hand, cytoskeletal proteins such as neurofilament light chain (NfL)^2^ and glial fibrillary acidic protein (GFAP)^3^ are currently under study for their ability to respectively capture neuroaxonal damage and astrocytic activation, but their dynamics are still poorly understood, likely due to long (weeks-to-months) half-life, especially in the case of NfL^4^.

Cell-free DNA (cfDNA) is a highly fragmented form of DNA that is shed in biological fluids as a consequence of apoptosis^5^, with an estimated half-life in blood of a few hours^6^. Applications of cfDNA profiling are already implemented in clinical practice and rely either on genetic information, as for prenatal diagnostics^7^ and tumor-specific mutations, or on disease-specific DNA methylation patterns, as in some types of cancers^8^. Specific DNA methylation patterns of a cell type of origin are preserved in cfDNA and recent studies have explored the possibility of using these methylation patterns in blood cfDNA to trace ongoing tissue-specific cell death throughout the organism^9^.

The CSF represents the most proximal and biologically relevant source of a “liquid biopsy” of the central nervous system (CNS). In fact, cfDNA has been detected in CSF from individuals with brain tumors^10^ as well as in healthy subjects, although in minute amounts (less than 1 ng)^11^. Moreover, CNS cells, particularly neurons, exhibit high levels of non-conventional DNA modifications, such as 5-hydroxymethylation (hmC), alongside common 5-methylation of cytosines (mC), both reflecting the transcriptional activity and regulatory.^12^ Hence, characterizing hmC and mC signatures of cfDNA originating from dying CNS cells could provide both a specific marker for neuronal damage but also critical molecular insights into disease mechanisms. However, sensitive detection of hmC remains technically challenging, with low cfDNA input representing a major limitation.

We here utilize duet evoC 6-base sequencing technology^13^ for simultaneous detection of mC and hmC in ultralow input of cfDNA extracted from CSF of people affected with various neurological conditions and healthy controls (HC). We aimed to assess the potential of CSF cfDNA to 1) enhance the sensitivity of detecting and quantifying CNS damage in real-time, and 2) dissect ongoing disease-specific pathological processes (**Fig. 1a**).

**Figure 1.**
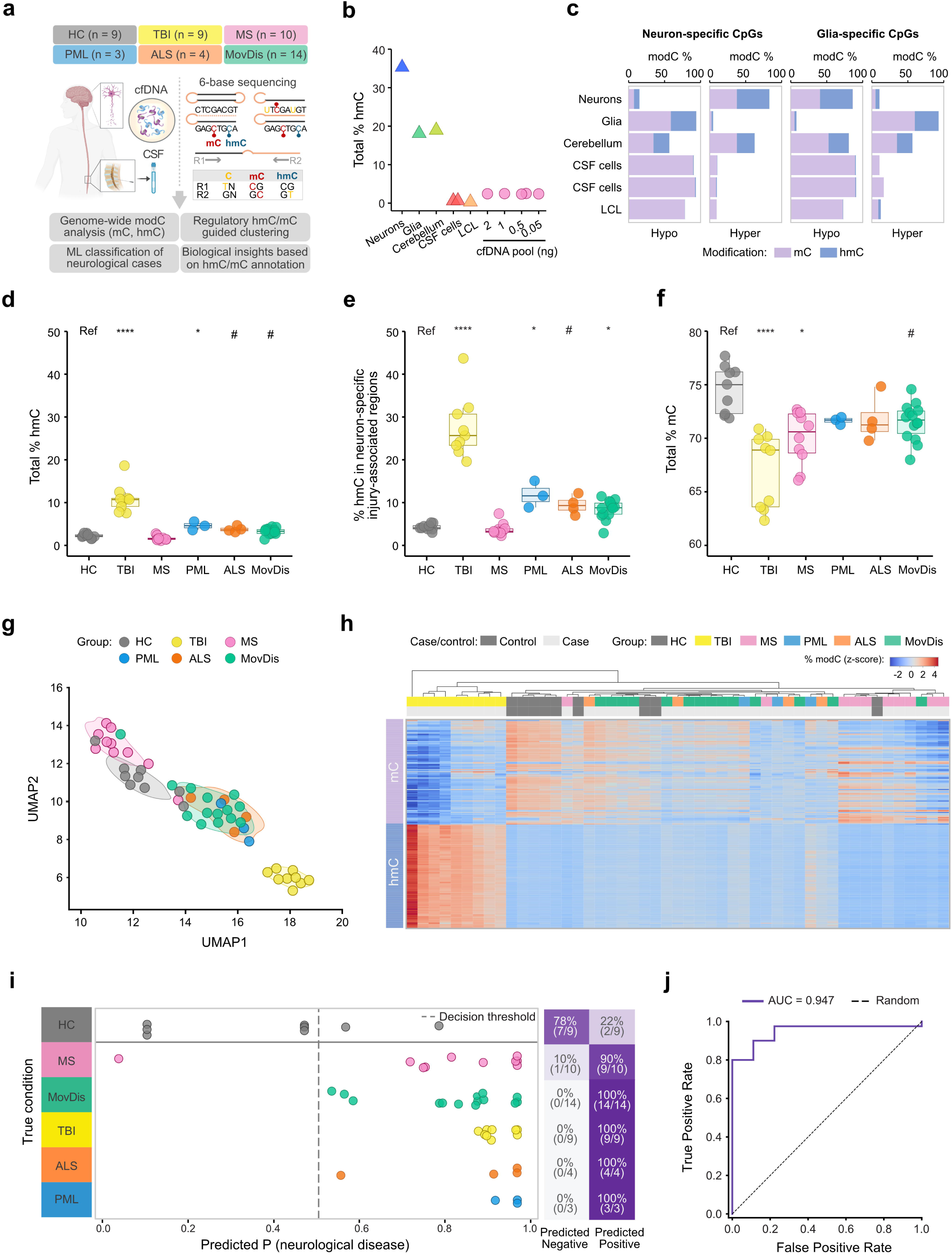
Simultaneous detection of hmC and mC in CSF cfDNA allows effective separation of neurological conditions from HC. **a,** Cohort overview and study design. HC: healthy control; TBI: traumatic brain injury; MS: multiple sclerosis; PML: progressive multifocal leukoencephalopaty; ALS: amyotrophic lateral sclerosis; MovDis: movement disorders. **b,** Mean total hmC percentage in control genomic (triangle) or CSF cfDNA (circles) samples. **c,** Contribution of hmC to total modC (mC+hmC) signal at CpGs hypo- or hypermethylated in neurons or glia, respectively. **d,** Mean total hmC percentage in individual CSF cfDNA samples of HC and various neurological conditions. **e**, Mean hmC percentage of neuron-specific injury-associated regions across individual CSF cfDNA sample groups. **f,** Mean total mC percentage across individual CSF cfDNA sample groups. Statistical significance in panels **d**, **e** and **f** is reported for pairwise comparison between neurological conditions and HC using Kruskal- Wallis test with post-hoc Dunn test and adjustment for multiple testing using Benjamini- Hochberg correction. #p<0.1, *p<0.05, **p<0.01, ***p<0.001, **** p<0.0001. **g,** UMAP of multi-omics factor analysis (MOFA) of hmC, mC, and modC across promoters and gene bodies. **h**, Heatmap representing hierarchical clustering based on hmC and mC mean signal in 100-chromatin states universal model, scaled by rows. The color gradient represents modification fraction Z-scores, row annotation for type of modification (hmC or mC), column annotation for sample classification. **i,** Dot plot of predicted probability of neurological disease for individual CSF cfDNA samples using leave-one-out cross- validation applied on hmC and mC signals in 100-chromatin state regions, with classification decision threshold set at 0.5 (dashed vertical line). Heatmap of prediction accuracy is reported on the right. **j**, Receiver operating characteristic curve for accurate identification of neurological disease with the developed model.

To ascertain the relevance of hmC detection for CNS cells, we first applied the evoC 6- base method to genomic DNA from sorted post-mortem neuronal (NeuN+) and glial (NeuN-) nuclei (**Suppl. Fig. 1a**), cerebellar tissue, CSF cells (composed of mainly lymphocytes) and the lymphoblastoid cell line NA12878 (LCL). Levels of total hmC (%) were, as expected, the highest in sorted neuronal DNA (35.3% at CpG sites), moderate in glia and mixed cerebellum tissue (18.1% and 18.9%, respectively), while almost absent (< 1%) in immune cells (**Fig. 1b**), confirming detection of neuronal DNA using hmC. Importantly, hmC contributed a striking 50% of the total modified cytosine (modC, i.e., mC + hmC) signal at neuron- and glia-specific CpG sites (**Fig. 1c**, **Suppl. Table 1**) derived using publicly available bisulfite conversion-based modC data (**Suppl. Fig. 1b-e**, see Methods). Given the ultra-low yield of cfDNA in CSF (<1 ng), the detection sensitivity of hmC and mC of 6-base sequencing method was tested using a dilution series (0.05, 0.5, 1 and 2 ng) of cfDNA extracted from pooled CSF samples. Duet evoC 6-base library preparation yielded successful cfDNA capture, with expected peak lengths (**Suppl. Fig. 2a**), high hmC/mC sensitivity and specificity, despite anticipated high duplication rate and limited coverage (**Suppl. Fig. 3**). The cfDNA samples exhibited noticeably higher total hmC content (mean 2.4%, range 2.3-2.5%) than immune cells (mean 0.5%, range 0.3- 0.6%), irrespective of the input (**Fig. 1b**). Thus, 6-base method proved to be not only feasible but, importantly, also sufficiently sensitive for hmC detection in cfDNA.

We next applied duet evoC 6-base sequencing to CSF cfDNA from a clinical cohort composed of severe traumatic brain injury (TBI, n = 9), multiple sclerosis (MS, n = 10), progressive multifocal leukoencephalopathy (PML, n = 3), amyotrophic lateral sclerosis (ALS, n = 4), movement disorders (MovDis, n = 14) and healthy controls (HC, n = 9) (cohort characteristics and sequencing QC measures are reported in **Suppl. Table 2** and **Suppl. Fig. 3,** respectively). Mean total hmC was significantly elevated in TBI (median 10.7%, range 7.6 - 18.6%, adjusted (adj. *p* < 0.0001) and PML (median 4.6%, range 3.6 - 5.6%, adj. *p* = 0.047) compared to HC (median 2.2%, range 1.5 - 2.9%), while a trend for higher hmC (adj. *p* < 0.1) was also observed in ALS and MovDis (**Fig. 1d**). Interestingly, hmC was highly correlated (r = 0.96, p < 0.0001) with non-CpG modC (**Suppl. Fig. 4a**), another epigenetic mark reported to be specifically enriched in neurons^12^. No statistically significant association of hmC levels with age or CSF NfL was observed (**Suppl. Fig. 4b-c**). Focusing on hmC at regions displaying differential hmC both in neurons compared to immune cells (“Neuron-specific”) and in TBI compared to HC samples (“Injury-associated”) (**Suppl. Fig. 4d**) further enriched hmC levels (mean increase 2.4-fold, range 1.6 – 2.9) and, importantly, enabled better separation of all neurological cases from healthy individuals, except for MS (**Fig. 1e**).

To refine neural signatures, we further integrated analyses of hmC with mC signals. Among control samples, sorted neurons, and to a lesser extent glia and cerebellum, displayed lower mean mC (respectively 43.1%, 62.4%, and 61.7%) compared to CSF cells (74.1-75.5%), directly reflecting the hmC contribution, while LCL exhibited both low mC (54.5%) and hmC (0.5%) content (**Suppl. Fig. 4e**). In the clinical cfDNA cohort, significantly lower total mC was observed both in TBI (median 68.9%, range 62.3 - 70.9%, adj. *p* <0.0001) and MS (median 70.6%, range 66.1 – 72.7%, adj. *p* = 0.012) compared to HC, which displayed the highest mC content (median 75.0%, range 71.9 – 77.7%) (**Fig. 1f**). Integrating hmC, mC, and modC levels at promoters and gene bodies using multi-omics factor analysis (MOFA) separated neurological conditions, especially TBI and MS, from HC (**Fig. 1g**). We then performed sample clustering and classification using mean methylation percentages in functionally relevant regions defined by the 100 chromatin-state annotations of the human genome from the International Human Epigenome Consortium (IHEC).^14^ Hierarchical clustering of mean hmC and mC levels across the regulatory features effectively clustered TBI and MS samples, while distinction between HC and other neurological diseases was only partial (**Fig. 1h**). To classify samples, we used machine learning approaches and trained a gradient-boosted tree model using a “leave-one-out” cross-validation method, where each sample served once as the test set and the remaining samples were used for model parameter selection and fitting (**Suppl. Fig. 5a**). Classification accuracy was 100% for all disease groups except MS and HC, which were correctly predicted in 90% and 78% of cases, respectively (**Fig. 1i**). This resulted in an area under the receiver operating characteristic curve (ROC-AUC) for detection of neurological cases *versus* HC of 0.947 (**Fig. 1j**). The strongest contribution to the model derived from the modifications listed in **Suppl. Fig. 5b-c**, including mC in regions annotated as brain enhancer (EnhA6_a).^15^ Altogether, these findings confirm the feasibility of 6-base sequencing in ultralow input CSF cfDNA and demonstrate the possibility of using hmC and mC modifications to detect neuronal damage across different neurological diseases.

Next, we sought to explore whether different epigenetic signatures across sample groups could inform about underlying disease biology. We selected 50,000 promoter or gene body regions that exhibited the highest variance for hmC, mC, and modC (for a total of 600,000 bins), and performed one-way ANOVA comparisons between each neurological condition and HC. TBI exhibited the highest number of significant (adjusted p < 0.05) bins compared to HC, followed by PML (**Suppl. Fig. 6a**). Comparing the top 2,000 bins, the majority were unique to each condition, with TBI having the largest proportion shared with other diseases (39.4%) (**Fig. 2a**, **Suppl. Fig. 6b**). In TBI, the signal was mostly driven by hmC in gene bodies, while MS had a more heterogeneous contribution across regions and type of modifications (**Suppl. Fig. 6c**). We next investigated whether the variant bins could inform cell-type contribution to the CSF cfDNA and conducted gene-set enrichment analysis (GSEA) for gene markers of relevant cell types. The analysis showed significant enrichment for immune cells in MS, neuronal and glial cells in TBI, and glial cells in MovDis compared to HC (**Fig. 2b-c** and **Suppl. Fig. 6d-g**). Given the lack of directionality in this variance-based analysis, we further explored cell-type composition of CSF cfDNA in each group, including HC, using deconvolution algorithms. Applying our internal genomic controls (neuron, glia and immune CSF cells) as deconvolution reference, TBI exhibited a large proportion of glia (0.39) and neurons (0.13), while MS had a higher contribution from immune CSF cells (0.92) (**Fig 2d**, top panel). To increase granularity, we used data from the Methylation Atlas^16^, which also estimated a numerically greater proportion from neurons, glia, and granulocytes, as well as a greater fraction of endothelial cells in TBI (adj. p = 0.009, **Fig. 2d** bottom panel and **Suppl. Fig. 7a**), and an enrichment for immune cells, especially T CD8+ cells (adj. p =0.009) in CSF cfDNA from MS (**Fig. 2d** bottom panel and **Suppl. Fig. 7b**), confirming the potential of hmC/mC analysis of CSF cfDNA to detect ongoing cell-type-specific cell death in the CNS.

**Figure 2.**
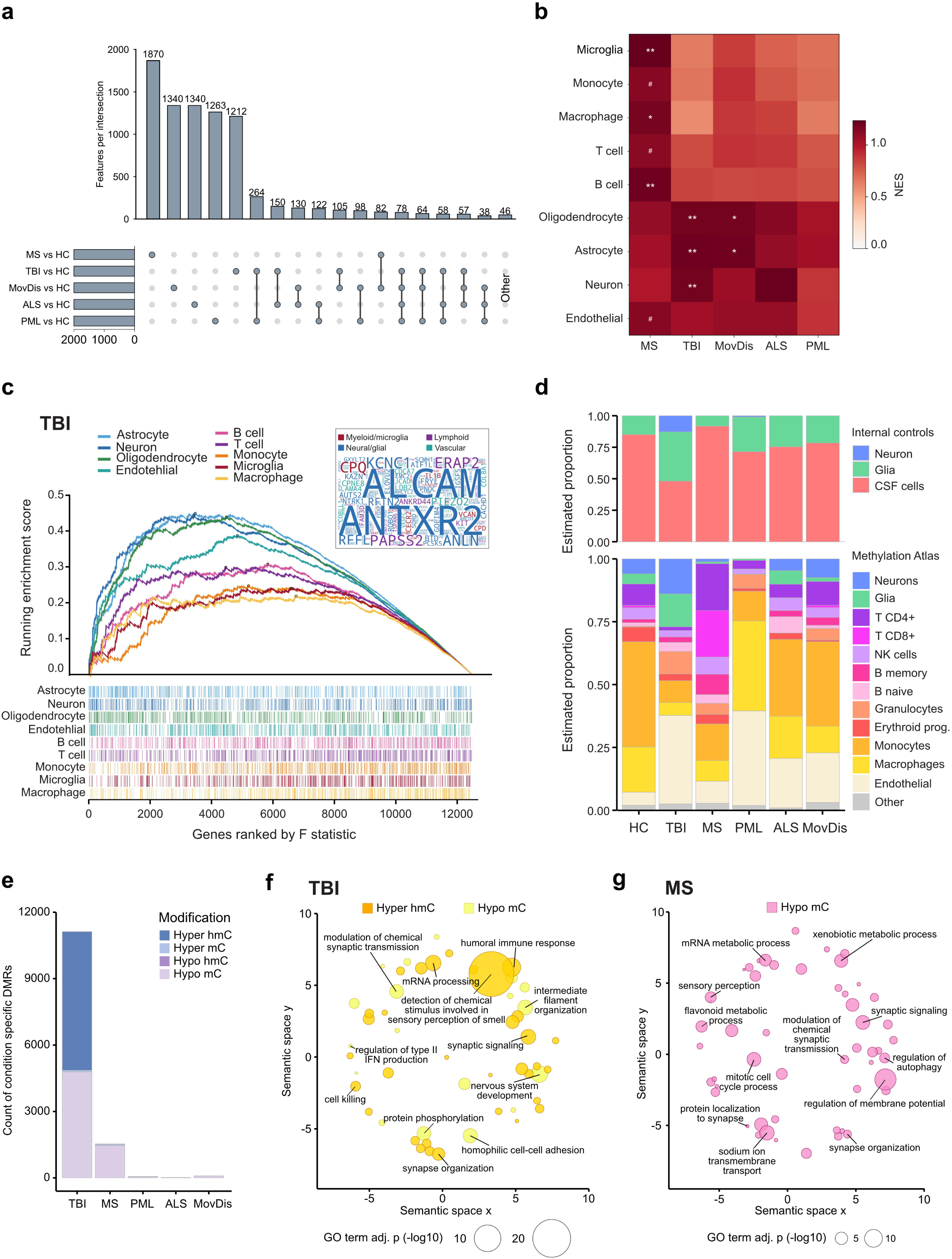
Differential (hydroxy)methylation of CSF cfDNA between neurological diseases and HC informs on disease-specific mechanisms in the CNS. **a,** Upset plot of the top 2,000 most variable bins in the comparisons between disease groups and HC, indicating numbers of condition-unique and shared features. **b**, Heatmap representing normalized enrichment scores (NES) from GSEA for cell-type specific gene set (CellMarker) applied on features ranked by increasing ANOVA F value for each comparison between neurological condition and HC. ^#^adj. p < 0.1, *adj. p < 0.05, **adj. p < 0.01. **c**, GSEA enrichment plot for cell-type specific ranked genes in the TBI *versus* HC comparison. The leading-edge gene cloud is reported in the top-right panel, color-coded by cell-type family. **d**, Stacked bar plot of mean estimated proportion of genomic DNA control samples (top) or Methylation Atlas cell types (bottom) across cfDNA sample groups. **e**, Stacked bar plot of condition-specific DMRs across disease groups compared to HC, color-coded by type of modification. **f-g**, GO BP terms enriched in hyper-hmC and hypo-mC TBI (**f**) and in hypo-mC MS (**g**) DMR, as summarized by semantic similarity by REVIGO.

We next investigated whether differentially methylated regions (DMRs) between disease and HC could inform on biological processes impacted by neurological pathology. Focusing on gene body regions that allowed for clear annotation and sufficient coverage (**Suppl. Table 3**), TBI had the highest number of condition-specific DMRs compared to HC (n = 11,125), of which majority displayed hyper-hmC (n = 6,265) or hypo-mC (n = 4,785), followed by MS (n = 1,539), where most DMRs were hypo-mC (n = 1,466) (**Fig. 2e**). Over- representation analysis of Gene Ontology Biological Process (GO BP) terms on these 3 categories of DMRs (**Suppl. Table 4**), summarized by semantic similarity with REVIGO^17^ (**Suppl. Tables 5-7**), identified in TBI enrichment of terms related to olfactory sensory neurons (top term “Detection of chemical stimulus involved in sensory perception of smell”), possibly related to the trauma dynamics in TBI, alongside broader CNS terms (e.g. “nervous system development”), and some immune terms (e.g. “humoral immune response”) (**Fig. 2f**). In hypo-mC DMRs in MS, interestingly, multiple terms referred to synaptic processes, including synaptic organization, protein localization, and signaling/transmission, possibly reflecting synaptic pathology in MS (**Fig. 2g**).

Taken together, simultaneous profiling of CSF cfDNA modifications from neurological conditions represents a novel sensitive method for detection of neuronal damage, enabling detection and investigation of neurological conditions in real-time.

## SUPPLEMENTARY MATERIALS

METHODS, including disclosures and data availability statement

## SUPPLEMENTARY TABLES

**Supplementary table 1. Neuron/glia-specific methylation sites.**

**Supplementary table 2. Demographic characteristics of CSF cfDNA clinical cohort.**

**Supplementary table 3. List of DMRs for hmC and mC annotated to gene bodies in neurological conditions compared to HC.**

**Supplementary table 4. List of GO terms enriched in TBI hyperhmC and hypomC, and MS hypomC DMRs.**

**Supplementary table 5. List of GO BP terms enriched in TBI hyper hmC DMRs as summarized by REVIGO.**

**Supplementary table 6. List of GO BP terms enriched in TBI hypo mC DMRs as summarized by REVIGO.**

**Supplementary table 7. List of GO BP terms enriched in MS hypo mC DMRs as summarized by REVIGO.**

## SUPPLEMENTARY FIGURES

**Supplementary Figure 1.**
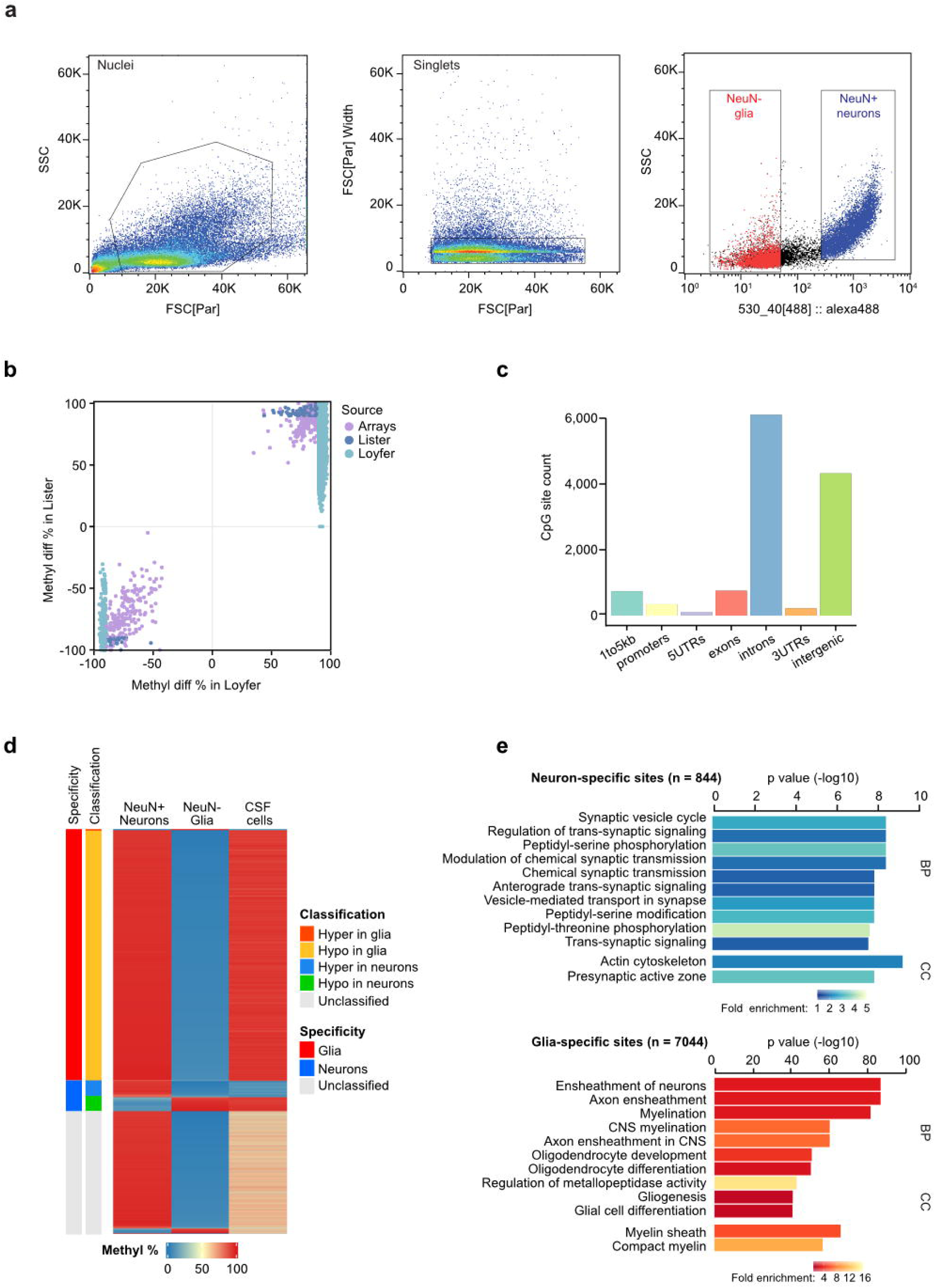
Neuron/glia-specific methylation sites. **a,** Sorting of NeuN+ (Neurons) and NeuN- (Glia) nuclei from snap-frozen brain tissue from control subjects for genomic control samples, defined by non-neurological cause of death. **b,** Scatter plot of delta (NeuN+ minus NeuN-) methylation percentage in the selected sites across the difference source used, showing high concordance between datasets. **c,** Genomic feature annotations of selected CpG sites. **d,** Heatmap showing methylation percentage in the selected CpG sites across NeuN+ and NeuN- samples from original source datasets, as well as in CSF cells. Site specificity (annotation on the left) was assigned to either neuron or glia when the signal differed not only from the other CNS cell type but also from CSF cells. (**e**) Horizontal bar plot depicting top 12 GO BP and Cell Component (CC) terms when performing functional enrichment analysis on neuron (top) and glia (bottom) specific sites with rGREAT package.

**Supplementary Figure 2.**
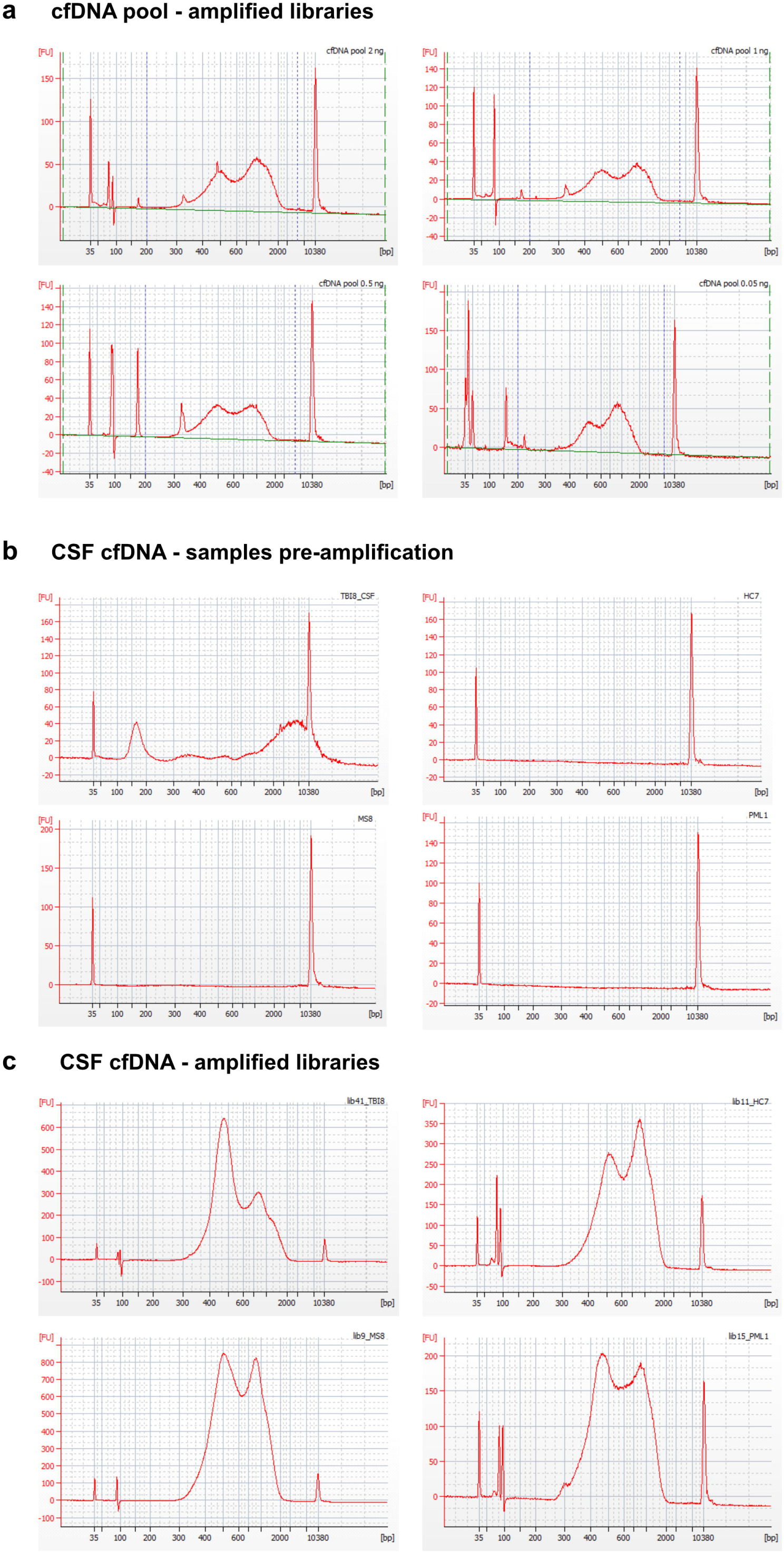
Duet evoC kit yields successful libraries for ultralow input of cfDNA CSF. **a**, Bioanalyzer electropherograms of libraries prepared with duet evoC kit from a pool of CSF cfDNA, starting from an input of 2, 1, 0.5 or 0.05 ng of DNA, showing fragment peaks at expected length (∼300 for mononucleosomal and ∼500 bp for dinucleosomal fragments). **b,** Examples of Bioanalyzer electropherograms of original samples before amplification, showing detectable DNA in TBI (top left panel), but not in HC (top right), MS (bottom left) and PML (bottom right). **c,** Examples of Bioanalyzer electropherograms of libraries prepared with duet evoC kit, after amplification, with fragment peaks at expected lengths. Same samples as in panel **b** are shown.

**Supplementary Figure 3.**
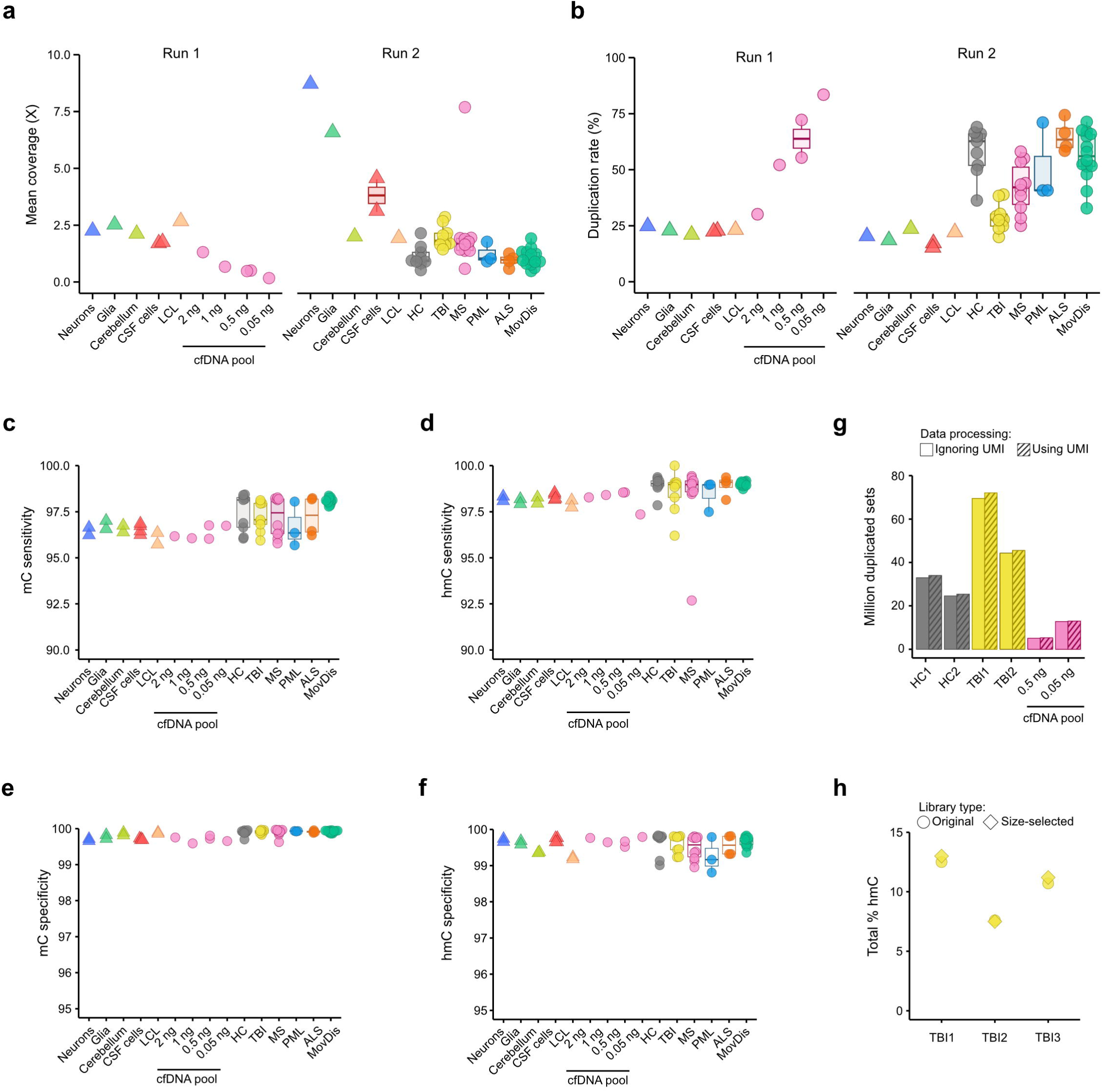
Quality control measures of CSF cfDNA libraries. **a-b,** Mean genome coverage (**a**) and duplication rate (**b**) for libraries prepared from control samples (Run 1) and clinical cohort as well as resequencing of genomic control samples (Run 2). **c-f,** QC measures for (hydroxy)methylation detection accuracy: sensitivity for mC (**c**) and hmC (**d**), target sensitivity >95%, and specificity for mC (**e**) and hmC (**f**), target specificity >99%. **g,** Barplot depicting number of duplicated sets (in millions) in libraries constructed with UMIs, detected either ignoring (no pattern) or using (striped pattern) UMIs for deduplication. **h,** Dotplot depicting total mean hmC (%) of whole genome in TBI samples before (original samples, circles) or after (size-selected, diamonds) size-selection for shorter fragments, indicating negligible interference of genomic DNA contamination on the analysis.

**Supplementary Figure 4.**
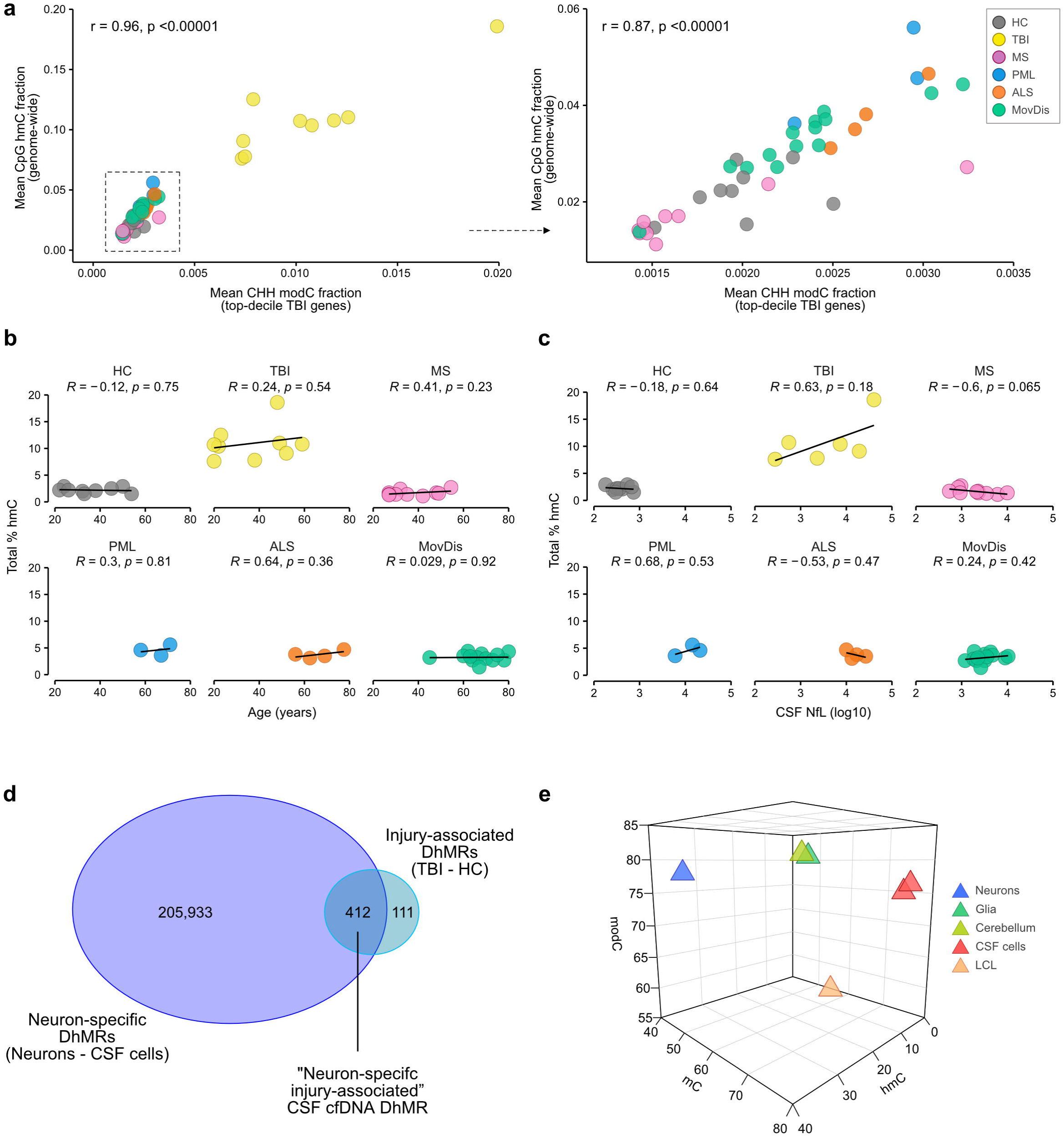
Hydroxymethylation correlation with molecular and clinical characteristics. **a,** Scatter plot of mean nonCpG methylation (CHH, x axis) and mean total CpG hydroxymethylation, including (left panel) or excluding (right) TBI samples, color-coded by disease group, showing high correlation between two epigenetic markers reported enriched in neurons. **b-c,** Scatterplot and linear fit lines correlating mean total hmC (%) with age (b) and CSF NfL (log10-transformed, c) in different disease groups. Pearson’s correlation coefficient and p value are reported. **d,** Venn diagram illustrating the selection method for neuron-specific injury-associated regions, derived from the intersection of the DhMRs between neurons and CSF immune cells (“Neuron-specific”, filtered for q value ≤ 0.1 and methylation difference ≥ 0.4) and between TBI and HC samples (“Injury-associated”, filtered for q value ≤ 0.1 and methylation difference ≥ 0.25). **e,** 3D scatterplot reporting mean total hmC (x axis), mC (y axis) and modC (hmC+mC, z axis) in genomic control samples.

**Supplementary Figure 5.**
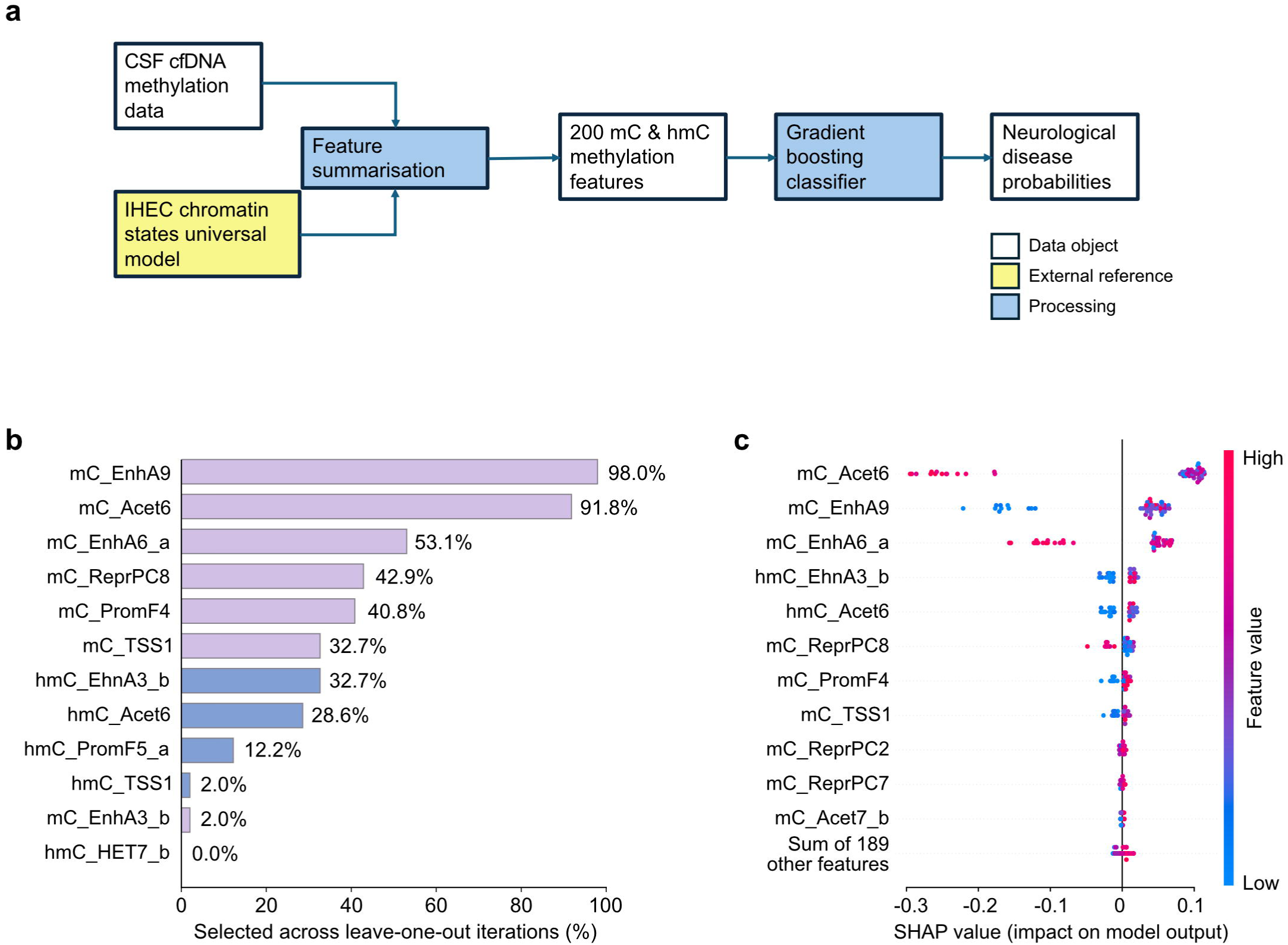
Classifier for neurological conditions using gradient-boosting model with “leave-one-out” validation. **a,** Overview of machine-learning algorithm used for sample classification. **b,** Horizontal bar plot of top 12 most informative features (mC or hmC) in the 100-chromatin state annotation model. A selection frequency of 100 indicates that the selected feature is always selected as informative in the iterations of the model. **c**, Scatter plot indicating SHAP values for the top 11 and the sum of the remaining features. SHAP values describe the relative importance and direction of individual features contribution to the model predictions. For example, high values of mC_Acet6 contribute negatively to the model probability (i.e. decrease the likelihood of a positive call).

**Supplementary Figure 6.**
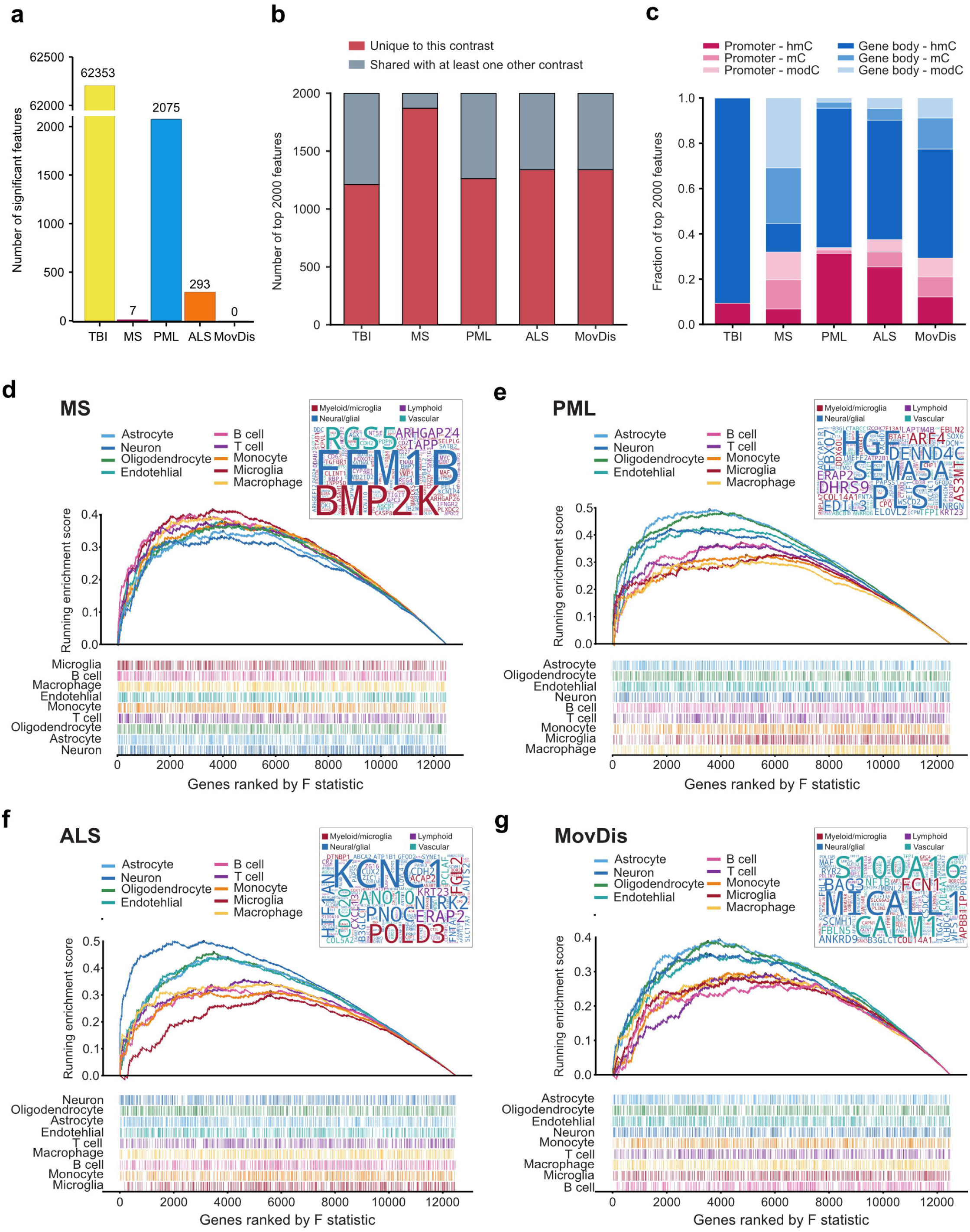
Cell-type enrichment using GSEA in different neurological conditions compared to HC. **a**, Barplot of significant (adj. p < 0.05) bins in each neurological condition compared to HC by ANOVA. **b**, Proportion of the top 2,000 bins that are unique to each comparison. **c**, Distribution of modification and region type in the top 2,000 bins per contrast. **d-g**, GSEA enrichment plots for cell-type specific genes in MS (**d**), PML (**e**), ALS (**f**), and MovDis (**g**) compared to HC, with leading-edge gene cloud reported in the top-right panel, color-coded by cell-type family.

**Supplementary Figure 7.**
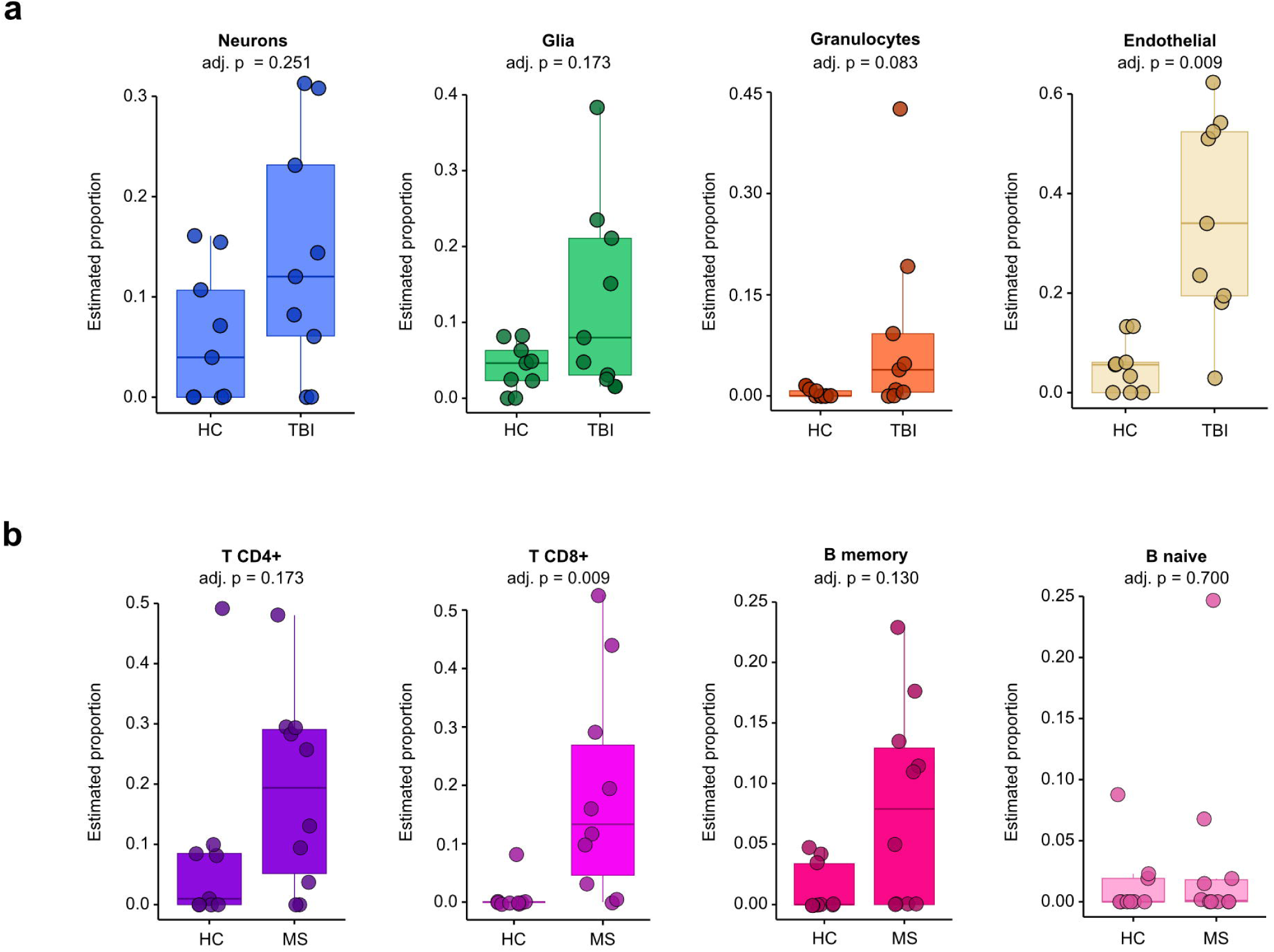
Cell-type deconvolution using the Methylation Atlas. Boxplots of estimated cell proportion of relevant cell types in TBI (**a**) and MS (**b**) compared to HC. Comparison by Wilcoxon test, corrected for multiple testing by Benjamini-Hochberg.

