## Supplementary material for "Advancing Neurological Liquid Biopsy via Six-Base Sequencing of Cerebrospinal Fluid Cell-Free DNA": Methods

### METHODS

#### Clinical cohort

CSF samples were collected from HC (n=9) and people with TBI (n=9), MS (n=10), PML (n=3), ALS (n=4), MovDis (n = 14). MovDis cases included 1 case of Parkinson's disease (PD), 8 progressive supranuclear palsy (PSP) and 5 multiple system atrophy (MSA).

CSF from patients with TBI was obtained through an external ventricular drain in the 48 hours following the trauma during their hospitalization in the neuro-critical care unit at Karolinska University Hospital (Stockholm). All patients had severe brain trauma, with Glasgow Coma Scale (GCS) between 3 and 8 upon admission. CSF for HC and non-TBI disease groups was obtained by lumbar puncture, performed during the diagnostic workup for neurological cases at Karolinska University Hospital (MS, ALS), Academic Specialist Centrum, Stockholm (MS, PD), and at Sahlgrenska Academy, Gothenburg (PSP and MSA). Clinical data were extracted from medical records by the treating physician. HC samples were collected from voluntary individuals with no history of neurological disease or familiarity for MS. PML cases were secondary to immunosuppressive therapy with rituximab in rheumatological diseases (lung sarcoidosis and rheumatoid arthritis).

Upon collection, CSF was centrifuged at 350 g (2000 g for TBI, MSA and PSP samples) and then stored at -80°C until further analysis.

#### Ethics

The study was conducted in line with the Declaration of Helsinki and ethical approval was obtained for each disease group from the regional ethical board in Stockholm (for HC, MS and PML STOPMS-II 2009/2107-31/2 and 2025-04961-02, for TBI 2005/1526-31/2, for ALS ALSrisc 2014/1815-31/4, 2018/1605-31 and 2021-06391-02, for PD 2020-03684) and in Gothenburg (for PSP and MSA, 460-13). Written informed consent was obtained by all participants before study inclusion (oral consent by next-of-kin in case of TBI samples).

#### Control samples

Cell-free DNA extracted from CSF from multiple individuals with MS (n = 90) was pooled in a single sample to reach measurable amount to use for technical validation.

Genomic DNA from relevant cell types was used as control. Neuronal (NeuN+) and glial (NeuN-) nuclei were extracted as previously described[1] from snap-frozen brain tissue from control subjects (defined by non-neurological cause of death) obtained from the Multiple Sclerosis and Parkinson's Tissue Bank, Imperial College London, approved by local ethical guidelines (**Suppl. Fig. 1a**). After tissue homogenization and nuclei extraction by ultracentrifugation in sucrose gradient, nuclei were labelled with Alexa Fluor 488 anti-NeuN antibodies (Invitrogen) and sorted by fluorescence activated cell sorting on a MoFlo™ high speed cell sorter.

CSF cells were extracted from one person with MS whose CSF was collected at baseline and during anti-CD20 treatment. Cells were collected by centrifugation at 440 g for 10 minutes, and immediately stored at -80°C after removal of the supernatant.

Cerebellum human tissue and NA12878 lymphoblastoid cell line genomic DNA were acquired pre-extracted from suppliers (Biochain and Coriell institute, respectively).

Genomic DNA was isolated from cells or nuclei using QIAmp DNA micro kit (Qiagen) following manufacturer's instructions, and then mechanically fragmented to 250-300 bp with Covaris M220 Focused ultrasonicator before library preparation following the manufacturer's instructions/parameters.

**Cell-free DNA extraction**

Cell-free DNA was extracted from 1 ml (TBI) or 4 ml (non-TBI) of cell-free CSF using QIAmp MinElute ccfDNA kit (Qiagen) following manufacturer's protocol. Briefly, samples were thawed and centrifuged at 16000 g for 10 minutes at 4°C to remove potential cell debris. The supernatants were mixed with an appropriate amount of Magnetic Bead Suspension, Proteinase K and Bead Binding Buffer and incubated for 10 minutes at room temperature (RT) in a HulaMixer (Invitrogen). DNA-loaded beads were isolated after 1-minute incubation in DynaMag™-15 magnet (Thermofisher), and DNA recovered with 200 µl Bead Elution Buffer. Beads were eliminated by 1-minute incubation in DynaMag™-2 Magnet (Thermofisher), and eluted DNA was mixed with 300 µl ACB buffer and loaded on a QIAamp UCP MinElute Column for purification. The column was centrifuged for 1 minute at 6000 g, loaded with 500 µl of ACW2 buffer after discarding the flow-through, centrifuged again at 6000 g for 1 minute, followed by another centrifugation at 20000 g for 3 minutes in absence of any buffer. The column filter was dried at 56°C for 3 minutes with the lid open, and the DNA was finally eluted in 50 µl of nuclease-free water by incubation at RT for 3 minutes, followed by centrifugation for 1 minute at 20000 g. The elution step was repeated twice (using the eluate from the first round) to increase the final DNA yield. DNA was quantified by Qubit dsDNA HS Assay kit (Invitrogen) and Bioanalyzer DNA High Sensitivity kit (Agilent), yielding measurable amounts only for TBI samples. For the remaining samples, the whole sample volume was concentrated down to 24 µl for library preparation using vacuum centrifugation with SpeedVac (Eppendorf) at RT for 20-25 minutes.

**Duet evoC library preparation and sequencing**

For control samples, 10 ng genomic DNA and 2 ng, 1 ng, 0.5 ng and 0.05 ng pooled cfDNA were used for library preparation. For TBI cfDNA samples, 1-2 ng were used for samples with low or high degree of long fragment contamination, respectively. Due to non-measurable DNA amount for non-TBI cfDNA samples, the whole sample volume was used, with an estimated total input of 100 pg. Libraries were prepared as per manufacturer's instructions. Lambda (methylated) and pUC19 (unmethylated) plasmids provided with the duet evo-C kit were spiked in each sample with the following dilution factors: 1:10 for 10 ng sample input, 1:100 for 1-2 ng sample input, and 1:1000 for lower starting input. The final amplification step included 7 PCR cycles for 10 ng starting input,

11 cycles for 1-2 ng starting input, and 13 cycles for input ≤ 500 pg. Libraries for control samples were first sequenced on Illumina NovaSeq 6000 SP-300 2x150bp (**Suppl. Fig. 3a-b** "Run 1"). Clinical cohort libraries, as well as genomic samples libraries, were sequenced on four lanes on Illumina NovaSeq XPlus 10B-300 with 2x150bp reads and 10% PhiX. The lowest concentrated cfDNA pool samples and some individual cfDNA samples (HC and TBI) were prepared with UMIs to verify whether duplication rate could be due to biological reasons, but little difference was detected upon de-duplication (**Suppl. Fig. 3g**). For three TBI samples, libraries were also prepared after short-fragment size selection using SPRI beads at 0.7x concentration, but little difference was detected in comparison with libraries prepared with the original sample (**Suppl. Fig. 3h**).

FASTQ files were processed using biomodal's duet data pipeline (version 1.6.0) which provides resolved genetic and epigenetic information in a single software workflow[2].

**Neuronal-specific CpG site reference compilation**

Four publicly available DNA methylation datasets of sorted NeuN+ and NeuN- nuclei from non-neurological controls were used as reference, two of which from whole-genome bisulfite sequencing (WGBS) studies (Lister et al. 2013[3], n = 2 paired NeuN+ and NeuN-, and Loyfer et al. 2023[4], n = 7 NeuN+ and 4 NeuN-) and two from studies conducted with Illumina Infinium HumanMethylation450K array (Guintivano et al. 2013[5], n = 29 and Gasparoni et al. 2018[6], n = 16, both with paired NeuN+ and NeuN-). After merging reads from CpG pairs and filtering for 10X coverage in WGBS data, we selected sites that displayed extreme methylation differences (≥ 90% in sequencing data and ≥ 70% in array data) between NeuN+ and NeuN-, so that the identified CpG sites were, for example, hypermethylated in neurons and hypomethylated in glia or vice versa. With this strategy, we identified 11323 CpG sites, which were cross validated between datasets (**Suppl. Fig. 1b, Suppl. Table 1**). These sites were mainly derived from introns and intergenic regions (**Suppl. Fig. 1c**). To determine neuronal- or glial-specificity, we then contrasted methylation in these sites with a whole-genome methylation dataset of CSF cells from people with MS (n = 13) available in-house, assigning site specificity to either neuron or glia when the signal differed not only from the other CNS cell type but also from CSF cells (**Suppl. Fig. 1d**). CSF cells were selected as contrast for CNS-cell methylation hypothesizing that DNA from CSF cells would be the most likely contributor to CSF cfDNA pool background. Conducting pathway analysis with the GREAT tool[7] highlighted enrichment for genes related to synaptic structures in case of neuron-specific sites, and for myelination and glial development in case of glia-specific sites (**Suppl. Fig. 1e**).

**Differential methylation analysis**

Calling of differentially methylated region (DMR) was performed using the modality XPLR analysis software (version 1.0.0). To enrich the 5-hydroxymethylcytosine (hmC) signal in regions that were specific to neurons and neuronal damage, DhMRs were called between genomic DNA from Neurons *versus* CSF cells (Neuron-specific) filtering for q value ≤ 0.01 and methylation difference ≥ 0.4, and between CSF cfDNA from TBI *versus* HC (Injury-associated) filtering for q value ≤ 0.01 and methylation difference ≥ 0.25. The regions shared between these contrasts were identified as "Neuron-specific injury-associated" regions (**Suppl. Fig. 4d**, **Fig. 1e**). Further, DMRs for hmC and 5-methylcytosine (mC) were also identified between neurological conditions and HC. DMRs for mC and hmC were called separately, genome-wide, using 5 kb windows, as well as on specific genomic annotations (gene bodies and promoters) using GENCODE gene annotations (release 44, GRCh38).

**Multi-Omics Factor Analysis (MOFA)**

Six bin-level categories were defined from GENECODE v44 annotation based on genomic feature (promoter, TSS ± 2kb; and gene body, full gene span) and modification (mC, hmC, and modC) contexts, respectively. Bins were retained if their summed denominator (pooling CG, CHG, and CHH reads) was ≥1 in at least 50% of samples, and the resulting fraction was converted to an M-value. For each category, the 50,000 most-variable bins across all samples were retained, giving up to ~300,000 bins in total and integrated using the Multi-Omics Factor Analysis (MOFA+, mofapy2)[8,9], which was fitted with view scaling enabled, ten factors, and a fixed random seed for reproducibility. The group structure was subsequently assessed using the learned factor matrix and group separation was quantified in MOFA factor space, using pairwise centroid distances and PERMANOVA. For visualization, factors were embedded with UMAP, with per-group kernel density estimates overlaid on the embedding.

**Classification of CSF cfDNA using universal chromatin state methylation features**

To classify CSF cell-free DNA samples as neurological disease (positive: TBI, MS, PML, ALS, and MovDis) versus healthy control (negative: HC), we derived methylation features from a 100-state chromatin annotation of the human genome.[10,11] For each sample, mean mC and hmC fractions were computed across CpG positions overlapping each of the 100 chromatin states, yielding a feature matrix of 200 features per sample (100 states × 2 modifications).

Classification was performed using a histogram-based gradient boosting classifier (scikit-learn HistGradientBoostingClassifier, class_weight='balanced'). Features were standardised within each cross-validation fold using zero-mean unit-variance scaling. Hyperparameters, including the number of boosting iterations, learning rate, and maximum tree depth, were tuned via an inner 3-fold cross-validation grid search, with the configuration maximising inner-fold balanced accuracy selected. The model was assessed using leave-one-out cross-validation (LOO-CV) as the outer loop, in which each sample served once as the test set and the remaining samples were used for inner hyperparameter selection and model fitting. Per-sample predicted probabilities from each LOO fold were aggregated to construct a single ROC curve and compute overall AUC, balanced accuracy, sensitivity, and specificity. Seed sensitivity was calculated by repeatedly running the classifier with 25 different starting seeds resulting in a mean AUC of 0.946 ± 0.012 [0.925, 0.975]. The model with AUC closest to the mean observed from the seed sensitivity test was chosen as the representative model. A permutation test permuting the cohort labels at random was conducted to generate the null distribution of model performance, after 5000 iterations of the model mean permuted AUC was 0.439 ± 0.165.

**Differential-bin ranking and cell-type enrichment**

For each pairwise comparison, the cohort was subset to the two groups compared (HC and each neurological disorder), and every bin (as defined for MOFA) was scored by a one-way ANOVA F-statistic (between-group over within-group variance) with Benjamini–Hochberg correction after median imputation of missing values. Because the F-statistic is a variance ratio it is direction-agnostic, ranking bins by how strongly group membership explains their variance without imposing a fold-change or effect-size threshold. Bins were mapped to genes (protein-coding only, divided in promoters or gene bodies), collapsed to one score per gene by maximum F, and used to rank the full gene list for competitive gene set enrichment analysis against CellMarker[12] cell-type signatures, restricted to a marker-only universe with recurrence-based consensus sets. Enrichment was assessed by a weighted running-enrichment statistic with a size-matched permutation null, and the leading-edge genes driving each significant cell type were displayed as per-cell-type gene clouds scaled by F and coloured by cell-type family.

**Cell-type deconvolution**

For cell-type deconvolution, mean mC in promoters and mean hmC in gene bodies was calculated for each of the CSF cfDNA samples, and the tissue reference samples, i.e. neurons, glia, and CSF cells. The CSF cfDNA samples were then deconvolved using non-negative least squares (NNLS, SciPy nnls) against the reference set of the tissue samples.

When using the Methylation Atlas reference[4], a custom subset was created using lymphocytic sub-types (naïve B-cells, memory B-cells, T-CD4, T-CD8, NK cells), as well as neuron, glial, endothelial and other immune cells as separate categories. The sample level methylation files were downloaded and the atlas created using the tools provided by the authors, and the "full" set of marker regions from the published methylation atlas. NNLS deconvolution (SciPy nnls) was performed on CSF cfDNA samples in our cohort using the sample mean modC of marker regions in the resulting atlas.

**DMR gene ontology analysis**

DMRs for TBI and MS were filtered for q value ≤ 0.01 and over-representation analysis for Gene Ontology (GO) Biological Process (BP), Cell Component (CC), and Molecular Function (MF) terms was conducted using goatools, restricting the analysis to terms including a minimum of 5 and a maximum of 500 genes. Term counts were propagated up the ontology, so that parent terms would get counts from child terms. P values were then corrected for multiple testing using Benjamini-Hochberg method, with FDR set at 0.05. For better visualization, the enriched GO-BP terms in the predominant condition-specific DMRs (hyper-hmC and hypo-mC for TBI, and hypo-mC for MS) were summarized by semantic similarity with REVIGO[13], using a cut-off value of 0.5 and the SimRel semantic similarity measure.

**Statistical analysis**

Demographic characteristics are summarized in **Suppl. Table 2** with median and range (minimum – maximum). Differences in global hmC and mC fraction across groups of CSF cfDNA samples were assessed by Kruskal-Wallis test, followed by post-hoc Dunn test between disease cases and controls. Resulting p values were adjusted for multiple comparisons by Benjamini-Hochberg correction. Cell-type deconvolution-based selected pairwise comparisons were analysed by Wilcoxon test, followed by Benjamini-Hochberg correction. Other statistical methods are reported with each analytical method description. Analyses were conducted in R v4.6.1 and Python 3.11.0.

### REFERENCES (Methods)

### DATA AVAILABILITY

Data are available from the lead contact upon reasonable request and are deposited at the https://ega-archive.org/ (EGAS50000002090). Full code notebooks will be found upon publication at https://gitlab.com/jagodiclab/ (separate deposition during review process). Additional information required to reanalyze the data is available from the lead contact upon request.


### ACKNOWLEDGEMENTS

We deeply appreciate and express our gratitude to patients and healthy donors who contributed with time and biological material to research, as well as the healthcare personnel that make the biomaterial collection possible.

We thank Dr. Annika van Vollenhoven for the help with FACS and the Center for Molecular Medicine Flow Cytometry Core for providing research facilities.

We thank the International Human Epigenome Consortium (IHEC) for providing access to reprocessed and harmonized epigenomic data from a broad collection of human cell and tissue types (https://ihec-epigenomes.org/epiatlas/data/).

The authors acknowledge support from the National Genomics Infrastructure in Stockholm funded by Science for Life Laboratory, the Knut and Alice Wallenberg Foundation and the Swedish Research Council, and SNIC/Uppsala Multidisciplinary Center for Advanced Computational Science for assistance with massively parallel sequencing. The computations were enabled by resources provided by the National Academic Infrastructure for Supercomputing in Sweden (NAISS), partially funded by the Swedish Research Council through grant agreement no. 2022-06725.

### FUNDING

This study received funding from Neuro Sweden, Neuro Stockholm, MS Forskningsfonden, the Swedish Brain Foundation, Swedish Research Council, and the Horizon Europe Framework Programme (project no.101136991). MJ's team was supported by the European Research Council (Epi4MS, grant agreement no. 818170), Knut and Alice Wallenberg Foundation, Swedish Research Council, Swedish Brain Foundation, Neurofonden, MS Forskningsfonden, StratNeuro, Stockholm County Council - ALF project and the Horizon Europe Framework Programme (project no. 101136991). KF has received research grants from Neuro Sweden, Neuro Stockholm, the Swedish Research Council, CIMED and ALF. PS is a Wallenberg Clinical Scholar and is also supported by the Swedish Research Council (grant no. 2024-02542) and the Region Stockholm (grant no.FoUI-1021688). FF is supported by the Swedish Research Council (grant no. 2023-02428) and the US Center for Disease Control and Prevention (grant no. R01TS000348). LN is upported by grants from Swedish State Support for Clinical Research (#ALFGBG-984087 and #ALFGBG-1005346), the Gothenburg Medical Society (#GLS-1001019), the MS Research Foundation, Neuro Sweden, the Research Foundation of the Multiple Sclerosis Society of Gothenburg (Neuro Gothenburg), Edit Jacobson Foundation, Amlövs Foundation, Novartis Foundation for Medical-Biological Research, unconditional research grants from Sanofi, Göteborg Foundation for Neurological

Research, Helena Ahlin's Foundation and Björnson Foundation. EPT was supported by Region Stockholm Clinical Research Appointment and ALF Funding, Karolinska Institutet, Swedish Research Council, Swedish Brain Foundation, and donations from Erik Gut and Family. FPiehl is supported by the Swedish Research Council (grant no. 2023-02639), the Region Stockholm (grant no. FoUI-987565), the Swedish Brain Fund (grant no. FO2025-0354) and Erling Perssons Foundation (grant no. 2024-0127).

### AUTHOR CONTRIBUTIONS

CSC, CRP, LK, MN, and MJ designed the study. CSC and CRP performed DNA extraction and library preparation. CN and MN performed sequencing data pre-processing. CSC compiled the neuronal-specific CpG site lists with the help of YH and performed methylation analysis at the genomic level. MS performed MOFA, variance analysis, and GSEA for cell type markers with the help of MN. CN performed functional annotation-based hierarchical clustering, machine learning classification, as well as D(h)MR analysis and cell-type deconvolution, with the help of AS, LS, TC, and FPuddu. MK managed sample biobanking. VDK, BE, KF, PS, CI, CS, FF, RC, LN, EPT, and FPiehl contributed to sample collection. CSC, LK, MN, and MJ wrote the manuscript with contributions from all authors. All authors read and approved the final manuscript.

### COMPETING INTERESTS

CN, AS, LS, TC, and FPuddu are current employees of biomodal limited and hold share options. KF has received research support from Merck, Novartis, Amicus and UCB, all outside the submitted work. PS has received lecture honoraria grants from AbbVie, MERZ, AstraZeneca, and fees for serving on DMC in clinical trials with Lundbeck. CI has consulted for Cytokinetics, Pfizer, BioArctic, Novartis, Tikomed, Ferrer, Amylyx, Prilenia, Mitsubishi and Abbvie. She is also a board member of Tobii Dynavox and of the Stiching TRICALS Foundation; all outside the submitted work. L.N. has received lecture honoraria from Biogen, Merck, Novartis, Teva and Sanofi, has served on advisory boards for ArgenX, Janssen, Merck and Sanofi, has received unconditional research grant from Novartis and Sanofi. FPiehl has received research grants from Denka, Janssen, Merck KGaA, Novartis, Pfizer and UCB, and fees for serving on DMC in clinical trials with Lundbeck and Roche, and preparation of expert witness statement for Novartis. CSC, CRP, MS, YH, MK, VDK, BE, CS, FF, RC, EPT, LK, MN, and MJ declare no competing interests.
